# Loss of NRMT1 allows expression of multiple differentiation pathways and alters transcription of secreted proteins in C2C12 myoblasts

**DOI:** 10.1101/2025.07.29.667488

**Authors:** John G. Tooley, Gao Zhou, Jack Forster, Cameron Jones, Frank Tedeschi, Christine E. Schaner Tooley

**Affiliations:** Department of Biochemistry, Jacobs School of Medicine and Biomedical Sciences, State University of New York at Buffalo, Buffalo, NY 14203, USA; Center for RNA Science and Therapeutics, Case Comprehensive Cancer Center, Case Western Reserve University, Cleveland, OH, 44106, USA

**Keywords:** NRMT1, methylation, satellite cells, differentiation, secretion

## Abstract

Muscle stem cells (satellite cells) retain their identity and function through expression of the paired homeobox transcription factor PAX7. PAX7 is able to both stimulate satellite cell proliferation and activate target genes involved in establishing myogenic identity, including myogenic factor 5 (MYF5) and the other myogenic regulatory factors (MRFs). Upregulation of the MRFs promotes commitment to the muscle lineage by initiating withdrawal from the cell cycle, upregulating expression of muscle-specific transcripts, and directing myoblast fusion. We have previously shown that knockout of the N-terminal methyltransferase NRMT1 in C2C12 mouse myoblasts results in significantly decreased *Pax7* expression, an inability of the cells to differentiate into myotubes, and abnormal upregulation of osteogenic markers. Here, we use RNA-sequencing to more comprehensively determine how loss of NRMT1 affects the transcriptional profile of proliferating and differentiating C2C12 myoblasts. We see that upon inducing differentiation, NRMT1 knockout cells can downregulate cell cycle, DNA replication, and histone gene expression. Though they also have significantly downregulated *Pax7* and *Myf5* expression, other muscle-specific transcripts are significantly increased over wild type, indicating the muscle transcriptional program is not completely inhibited. However, signaling pathways involved in the differentiation of other types of mesenchymal and hematopoietic lineages are also increased with NRMT1 loss and expression of chemotactic genes is downregulated. Together, these data indicate that NRMT1 knockout cells can upregulate genes needed for cell cycle withdrawal and muscle specification but fail to suppress markers of other lineages and activate normal chemotactic signaling, which may lead to the observed differentiation phenotypes.

## Introduction

Muscle growth and regeneration occur through activation of satellite cells, stem cells resident within skeletal muscle tissue. These satellite cells remain in a quiescent state until they receive a signal to initiate differentiation (Yin et al., 2013). Activated satellite cells divide asymmetrically to produce a new subset of satellite cells that withdraw from the cell cycle and return to quiescence and a subset of myogenic precursor cells (myoblasts) that proliferate, differentiate, and contribute to myofiber formation (Addicks et al., 2019; Kuang et al., 2007). The paired homeobox transcription factor PAX7 is responsible for maintaining satellite cell function and identity. It regulates satellite cell self-renewal and proliferation, as well as helps initiate the differentiation process through upregulation of the myogenic regulatory factor (MRF) myogenic factor 5 (MYF5) (Addicks et al., 2019; Günther et al., 2013). MYF5 and the other MRFs, myogenin (MYOG), myogenic differentiation 1 (MYOD), and myogenic factor six (MYF6/MRF4), are transcription factors that promote muscle differentiation through upregulation of muscle-specific transcripts, cell cycle withdrawal, and myotube/myofiber formation (Hernández-Hernández et al., 2017; Lamarche et al., 2021; Rajabi et al., 2014). Loss of PAX7 results in dysregulation of the MRFs, cell-cycle arrest, progressive loss of satellite cells, and severe muscle atrophy (Lepper et al., 2011; Seale et al., 2000; von Maltzahn et al., 2013).

We have previously shown that loss of N-terminal RCC1 methyltransferase 1 (NRMT1) results in significantly decreased *Pax7* expression in C2C12 mouse myoblasts (Tooley et al., 2021). NRMT1 is the first identified eukaryotic N-terminal methyltransferase (Tooley et al., 2010; Webb et al., 2010). It is a distributive N-terminal trimethylase that methylates a canonical X-Pro-Lys N-terminal consensus sequence and an extended non-canonical N-terminal consensus sequence (Petkowski et al., 2013; Petkowski et al., 2012). Biochemically, N-terminal methylation has primarily been shown to regulate protein-DNA interactions (Cai et al., 2014; Chen et al., 2007; Dai et al., 2013), but it has also been shown to regulate protein-protein interactions and protein stability (Faughn et al., 2018; Nevitt et al., 2018). Biologically, N-terminal methylation has been shown to play a role in stem cell maintenance (Catlin et al., 2021; Tooley et al., 2021). NRMT1 knockout (*Nrmt1^-/-^*) mice exhibit severe neurodegenerative phenotypes that result from misregulation of the neural stem cell (NSC) pools (Catlin et al., 2021). They exhibit an early depletion of NSCs from both niches and a corresponding increase in intermediate progenitor and neuroblast populations (Catlin et al., 2021). The neuroblasts appear to differentiate and migrate correctly, though they fail to exit the cell cycle and ultimately begin undergoing apoptosis by six weeks (Catlin et al., 2021).

In C2C12 myoblasts, CRISPR/CAS9-mediated loss of NRMT1 has an opposite effect. It decreases proliferation and inhibits myotube formation (Tooley et al., 2021). This corresponds with significantly decreased *Pax7* expression, and interestingly, abnormal upregulation of markers associated with osteoblast differentiation (Tooley et al., 2021). These data indicate that C2C12 myoblasts are unable to differentiate correctly with NRMT1 loss, though it remains unclear mechanistically how this is occurring. It is possible that loss of *Pax7* expression results in the failure to initiate the entire muscle-specific transcription program, which results in loss of myoblast identity and allows for transdifferentiation. The tumor suppressor retinoblastoma protein (RB) is a NRMT1 substrate (Tooley et al., 2010), and its phosphorylation status, overall stability, and ability to repress cell cycle target genes are altered upon NRMT1 loss (Catlin et al., 2021). Therefore, it is also possible that loss of NRMT1 could be disrupting myoblast differentiation through inhibiting cell cycle withdrawal, as upregulation of RB function is needed for growth arrest of differentiating myoblasts (Kitzmann & Fernandez, 2001). Finally, myosin light chain 9 (MYL9) is also a substrate of NRMT1, and loss of N-terminal methylation of MYL9 alters cytoskeletal rearrangements (Nevitt et al., 2018), indicating this could disrupt myoblast migration into myotubes.

To distinguish between these possibilities, we performed RNA-sequencing to comprehensively determine how loss of NRMT1 affects the transcriptional profile of proliferating and differentiating C2C12 myoblasts. First, we confirm that loss of NRMT1 significantly decreases *Pax7* expression both under basal growth conditions and after one day exposure to differentiation media as compared to control cells. Interestingly, we also see that some, but not all, of the MRFs are correspondingly downregulated, and there is little difference between the control and knockout cells in downstream muscle-specific transcripts, including actins, myosins, and members of the troponin complex, after one day of differentiation. We also see that NRMT1 knockout (KO) cells are able to downregulate cell cycle, DNA replication, and histone genes upon induction of differentiation, which corresponds to the previously observed decrease in proliferation (Tooley et al., 2021). We find the biggest differences between the NRMT1 KO cells and the wild type (WT) C2C12 cells are that after one day of differentiation, the KO cells upregulate genes involved in the differentiation of mesenchymal and hematopoietic lineages and downregulate genes involved in chemotaxis. These data indicate that, while NRMT1 does not completely block muscle cell differentiation, it does inhibit singular commitment to the muscle lineage and may also block the signaling necessary for myoblast migration and fusion into myotubes.

## Materials and Methods

### Cell culture and *in vivo* differentiation

Wild type C2C12 mouse myoblasts and previously generated C2C12 myoblasts with CRISPR/CAS9-mediated loss of NRMT1 (Tooley et al., 2021) were grown in Dulbecco’s Modified Eagle Medium (DMEM; ThermoFisher, Grand Island, NY) supplemented with 10% fetal bovine serum (FBS; Bio-Techne/Atlanta Biologicals, Minneapolis, MN) and 1% penicillin-streptomycin (P/S; ThermoFisher) at 37°C and 5% CO_2_ on tissue cultured-treated plastic (Corning, Corning, NY). Differentiation was induced by growing the cells in differentiation media (DMEM, 2% horse serum, 50 nM insulin (MilliporeSigma, Burlington, MA)) for 24 hours. 2 x 10^8^ cells per sample were harvested in lysis buffer (20 mM Tris-HCl (pH 7.4), 150 mM NaCl, 5 mM MgCl_2_, 100 μg/ml cycloheximide, 1 mM DTT), lysed with a syringe, and spun at 4°C at 14K RPM for 10 min. Super was flash frozen in liquid nitrogen and sent to the Case Western Reserve University Center for RNA Science and Therapeutics for processing.

### RNA isolation and sequencing

Total RNA was extracted using TRIzol reagent (ThermoFisher) following the manufacturer’s protocol with minor modifications. For liquid samples, 800 µL of TRIzol was added to 200 µL of sample and incubated at room temperature (RT) for 5 mins. 0.2 mL chloroform (MilliporeSigma) was added. Samples were shaken vigorously by hand for 30 seconds, followed by a 10 min incubation at RT. Phase separation was performed by centrifugation at 12,000 × g for 15 mins at 4°C. The upper aqueous phase was carefully removed, and the RNA was precipitated by adding 1 μL of glycoblue and 0.5 mL of isopropanol per 1 mL of TRIzol used, followed by a 10 min incubation at RT. Samples were centrifuged at 12,000 × g for 10 mins at 4°C. The RNA pellet was washed twice with 75% ethanol and centrifuged at 12,000 × g for 5 mins at 4°C after each wash. Pellets were air-dried briefly at RT and resuspended in nuclease-free water.

For RNA-sequencing (RNA-seq), RNA was further purified by the RNeasy kit (Qiagen, Venlo, Netherlands) according to the manufacturer’s instructions. RNA concentration was measured using a NanoDrop spectrophotometer (ThermoFisher). Samples were diluted to 50 ng/µL and 20 µL aliquots were prepared in 1.7 mL tubes for RNA-seq library construction. The CWRU Genomics Core generated RNA-seq libraries using the NEB Next Ultra II Directional RNAseq kit according to the manufacturer’s instructions. The libraries were sequenced by the CWRU Genomics Core on a single lane of the NovaSeq X using 75-cycle single-end sequencing. For verification of RNA-sequencing results with quantitative real-time PCR (qPCR) (**Sup. Fig. 1a-e**), 1 μg of RNA was converted into cDNA using the SuperScript First-Strand Synthesis System (ThermoFisher). qPCR was performed with 2x SYBR green Master Mix in a CFX96 Touch Real-Time PCR Detection System (Bio-Rad, Hercules, CA). Transcript expression levels were determined using the ΔΔCT quantification method. Mouse primer sequences were as follows: *Myf5* forward 5’- CCTGTCTGGTCCCGAAAGAAC-3’ and reverse 5’- GACGTGATCCGATCCACAATG-3’; *Pax7* forward 5’- CCGTGTTTCCCATGGTTGTG-3’ and reverse 5’- GAGCACTCGGCTAATCGAAC-3’; *Mmp9* forward 5’- CTTCTGGCGTGTGAGTTTCCA-3’ and reverse 5’- ACTGCACGGTTGAAGCAAAGA-3’; *Robo1* forward 5’- GAACCTGCCACTCTGAACTG-3’ and reverse 5’- GTCCATGGACTATGCGTAAG -3’; and *Fgf7* forward 5’- CTCTGCTCTACAGATCATGC-3’ and reverse 5’- CATAACTTCTGGTGTGTCGC-3’.

### Data analysis

Quality control analysis was done using FastQC (v0.11.8; RRID:SCR_014583). The RNA-sequencing reads were aligned using STAR v2.5.3a (RRID:SCR_004463). Hierarchical cluster analysis (HCA) to assess sample relationships was performed through DESeq2 and found that knockout sample #1 from day 1 was a significant outlier (**Sup. Fig. 1f,g**), so this sample was removed from data analysis. The isoform table for RNA was the input for DESeq2 analysis, which was also used to construct the volcano plots. GO analysis was performed using the Database for Annotation, Visualization, and Integrated Discovery (DAVID) from the National Institutes of Health (Huang da et al., 2009; Sherman et al., 2022). Interaction networks were made using the STRING: protein query function in Cytoscape 3.10.3 (Shannon et al., 2003). Heatmaps were made using GraphPad Prism 10.5.0.

## Results

### Expression patterns in C2C12 cells after one day of differentiation

For the RNA sequencing experiments, mRNA was harvested from wild type (WT) C2C12 cells and C2C12 cells with CRISPR/CAS9-mediaed knockout (KO) of NRMT1 (Tooley et al., 2021) during basal growth conditions (day 0) and after 24 hours of growth in differentiation media (day 1). When comparing transcripts expressed in WT cells at day 1 versus WT cells at day 0, we found that 224 genes were significantly upregulated greater than two-fold (**Fig. 1a**). Though *Pax7*, *Myf5*, and *MyoD* expression remained constant, *MyoG* expression was significantly upregulated, as was expression of motor proteins and genes involved in skeletal muscle contraction, including actins, myosins, and members of the troponin complex (**Fig. 1b-d**). *Mrf4* transcripts were not detected at either timepoint, which is consistent with its later expression in the differentiation process (Hernández-Hernández et al., 2017). These data confirm that *Pax7*, *Myf5*, and *MyoD* are already expressed in C2C12 myoblasts under normal growth conditions, and *MyoG* expression increases upon initiation of differentiation, as previously described (Andrés & Walsh, 1996).

**Fig. 1.**
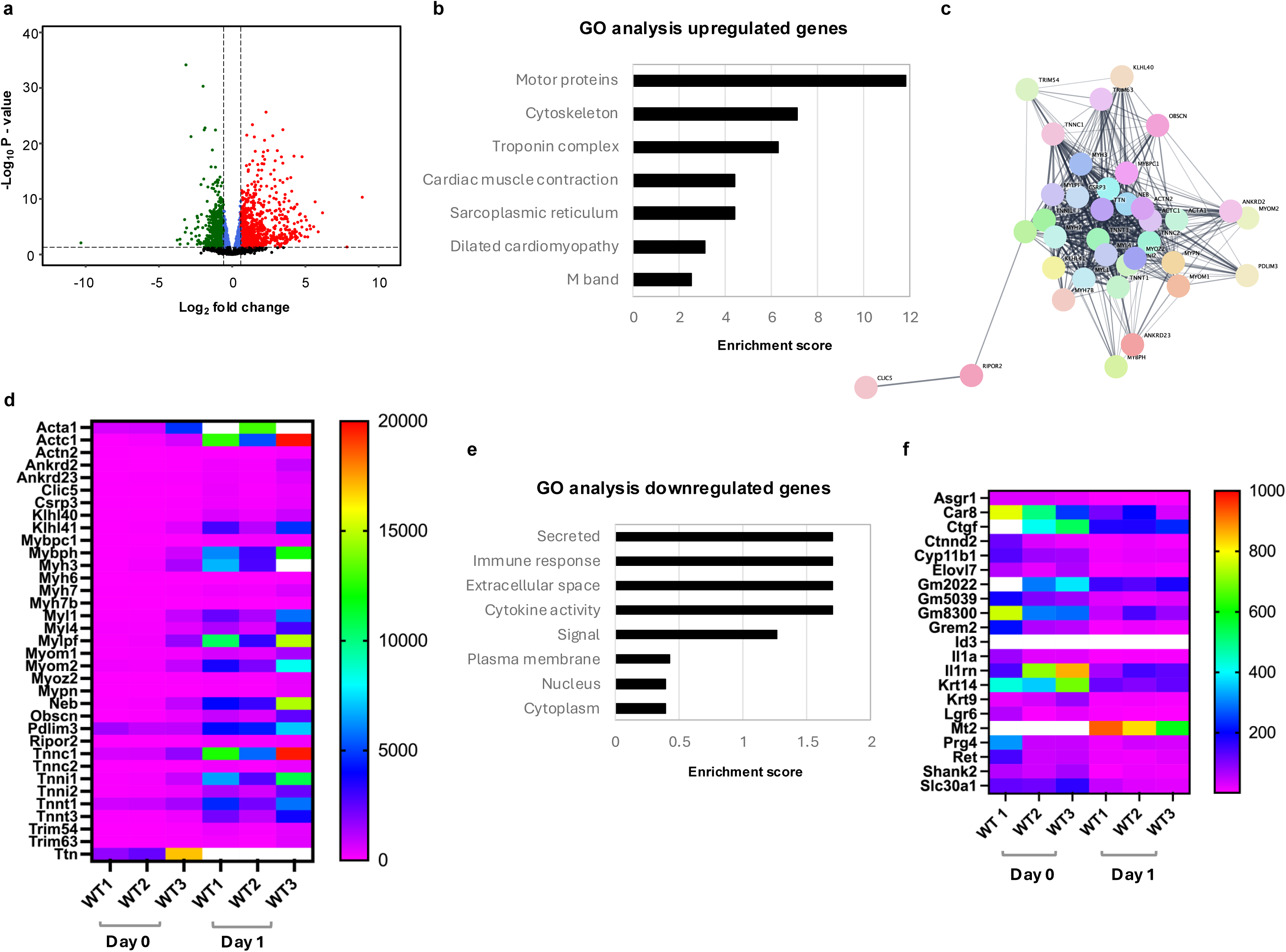
WT day 1 compared to WT day 0. **a** Volcano plot showing genes differentially expressed in wild type (WT) C2C12 cells between day 1 and day 0. Blue dots = p-value < 0.05 & Log_2_FC ≤ Log_2_(1.5); green dots = p-value < 0.05 & Log_2_FC < -Log_2_(1.5), red dots = p-value < 0.05 & Log_2_FC > Log_2_(1.5); black dots = p-value ≥ 0.05. **b** GO analysis of 224 genes upregulated greater than two-fold at day 1. **c,d** Interaction network and heatmap of muscle-specific transcripts upregulated at day 1. **e, f** GO analysis and heatmap of 21 genes downregulated greater than two-fold at day 1. Scale bars on heatmaps denote normalized counts. White bars indicate values are out of range.

In contrast, only 21 genes were significantly downregulated greater than two-fold after one day of differentiation (**Fig. 1a**), and these included genes that encode secreted and membrane proteins and genes involved in the immune response (**Fig. 1e,f**). Many of these genes are involved in differentiation outside of the muscle lineage. Solute carrier family 30a1 (SLC30a1) is involved in neural crest development (Xia et al., 2021). Asialoglycoprotein receptor 1 (ASGR1) is involved in immune cell differentiation, specifically the monocyte to macrophage transition (Shi et al., 2023). Connective tissue growth factor (CTGF) promotes both adipogenic and osteoblastic differentiation of mesenchymal cells (Battula et al., 2013; Yan et al., 2022). Interleukin 1α (IL1A) and interleukin 1 receptor antagonist (IL1RN) can also promote osteoblastic differentiation (Sarsenova et al., 2021; Zou et al., 2021). Leucine-rich repeat-containing G-protein-coupled receptor 6 (LGR6) plays a role in both muscle and bone differentiation (Khedgikar et al., 2022; Kitakaze et al., 2023), though it has been shown to decrease in expression one day after induction of differentiation in C2C12 cells (Kitakaze et al., 2023), consistent to what is seen here. Taken together, after one day exposure to differentiation media, muscle-specific transcripts increase expression in WT C2C12 cells, while transcripts of genes involved in the differentiation of other lineages decrease.

### Expression patterns in NRMT1 knockout C2C12 cells

When comparing expression patterns in the NRMT1 KO cells between day 1 and day 0, the first noticeable difference from WT cells is that *Pax7* and *Myf5* transcripts have negligible expression at both timepoints (**Sup. Fig. 1a,b**). However, more like WT, *MyoD* exhibits high, constant expression (data not shown), and *MyoG* expression significantly increases (**Fig. 2d**). In addition to *MyoG*, there were 474 genes significantly upregulated over two-fold after one day of differentiation (**Fig. 2a**). Also increasing were muscle-specific transcripts, including transcripts for proteins involved in cardiomyopathy, motor proteins, calmodulin binding proteins, and proteins in the troponin complex (**Fig. 2b-d**). These data indicate that even though NRMT1 KO cells exhibit low expression of *Pax7* and *Myf5*, they can maintain *MyoD* and *MyoG* expression and activate many of the same genes in the muscle transcriptional program as the WT cells. They also explain how NRMT1 KO is not lethal in myoblasts, as MYOG is the only MRF required for viability (Nabeshima et al., 1993).

**Fig. 2.**
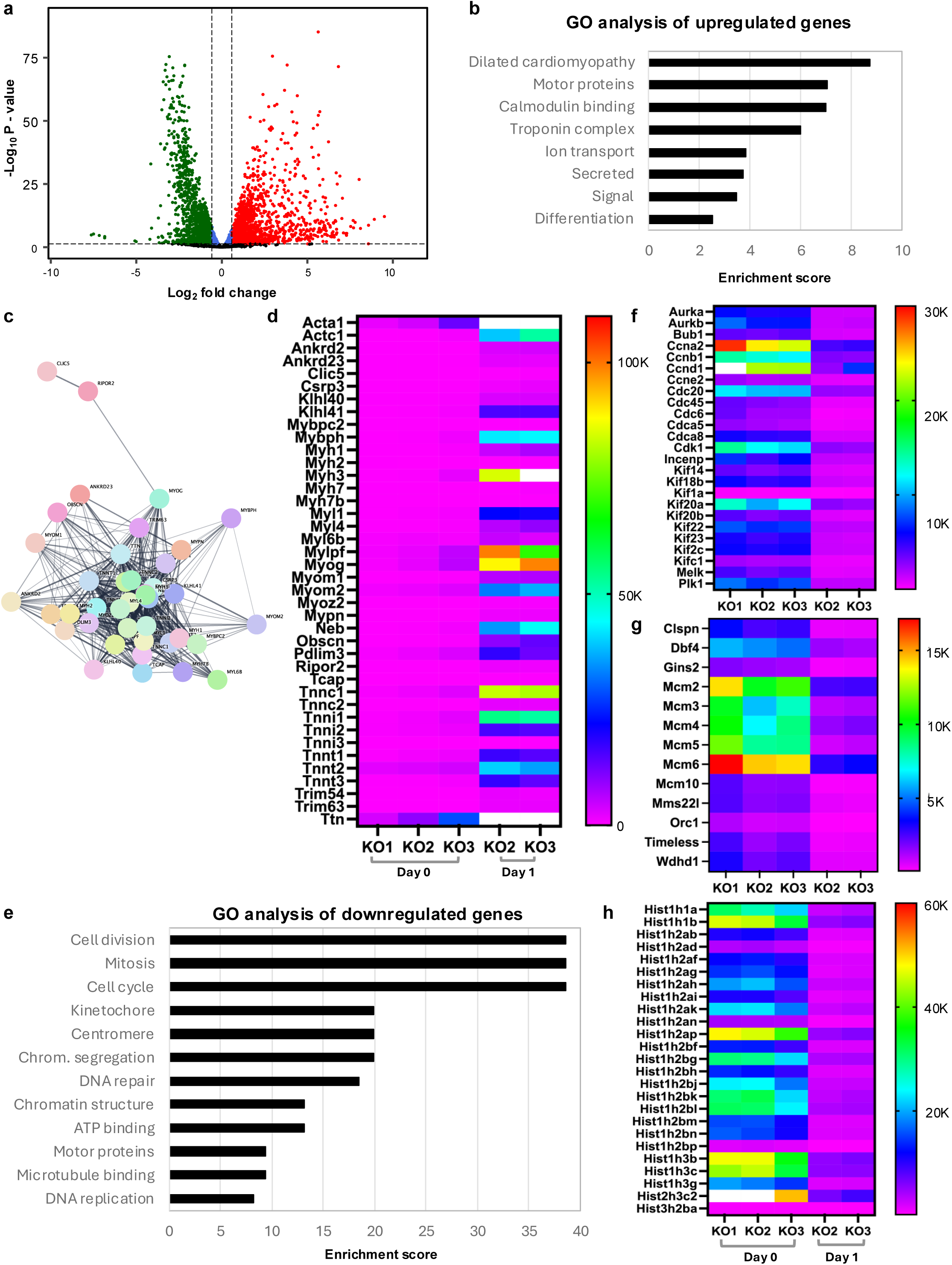
KO day 1 compared to KO day 0. **a** Volcano plot showing genes differentially expressed in NRMT1 knockout (KO) C2C12 cells between day 1 and day 0. Blue dots = p-value < 0.05 & Log_2_FC ≤ Log_2_ (1.5); green dots = p-value < 0.05 & Log_2_FC < -Log_2_(1.5), red dots = p-value < 0.05 & Log_2_FC > Log_2_(1.5); black dots = p-value ≥ 0.05. **b** GO analysis of the 474 genes upregulated greater than two-fold at day 1. **c.d** Interaction network and heatmap of muscle-specific transcripts upregulated at day 1. **e** GO analysis of the 356 genes downregulated greater than two-fold at day 1 and heatmaps of those genes that play a role in (**f**) cell division/cell cycle, (**g**) DNA replication, and (**h**) chromatin structure. Scale bars on heatmaps denote normalized counts. White bars indicate values are out of range.

In contrast to the 25 downregulated genes in WT cells, we found 356 genes were significantly downregulated greater than two-fold in KO cells after one day of differentiation (**Fig. 2a**). Also unlike WT cells, these genes were primarily involved in cell cycle, DNA repair, and chromatin structure (**Fig. 2e-h**). Downregulated cell cycle genes represent a variety of families including, the aurora kinases (AURKA and AURKB), the cyclins (CCNA2, CCNB1, CCND1, and CCNE2), a cyclin-dependent kinase (CDK1), members of the kinesin superfamily (KIFs), and a polo-like kinase (PLK1) (**Fig. 2f**). Downregulated genes involved in DNA replication include most of the mini-chromosome maintenance (MCM) proteins that make up the MCM2-7 complex (**Fig. 2g**), a DNA helicase necessary for unwinding the double helix (Lei, 2005). Interestingly, 52 of the 356 downregulated genes were histone proteins (**Fig. 2h**). Traditionally, histone protein expression is globally repressed either at the end of S phase, in response to a large amount of DNA damage, or at the initiation of differentiation (Mei et al., 2017; Stein et al., 1996; Su et al., 2004). Given that a large amount of cell cycle genes are also downregulated, and loss of NRMT1 decreases the growth rate of C2C12 cells (Tooley et al., 2021), these data indicate that these cells are withdrawing from the cell cycle in response to the initiation of differentiation.

### Differences in transcript expression during basal growth

When comparing KO cells at day 0 to WT cells at day 0, we found that 101 genes were significantly upregulated greater than two-fold in the KO cells (**Fig. 3a**). While *MyoD* and *MyoG*, were slightly upregulated, they did not make the two-fold cut off (data not shown). Genes that did make the cut off included those that encode for secreted proteins/proteins with signal peptides and proteins involved in interferon (INF) signaling, including many guanylate-binding proteins (GBPs) (**Figs. 3b,c**). INF-γ stimulation has been shown to inhibit myogenesis by reducing transcription of muscle-specific genes, including *MyoG* (Londhe & Davie, 2011). This is accomplished through activation of the major histocompatibility complex class II transactivator CIITA, which can recruit histone deacetylases and other co-repressors (Xu et al., 2008). While transient INF-γ signaling has been shown to be beneficial for muscle healing and regeneration (Cheng et al., 2008; Foster et al., 2003), exposure to exogenous INF-γ decreases the proliferation and myotube formation of myoblasts in culture (Kalovidouris et al., 1993; Tomita & Hasegawa, 1984). It is thought that this is through CTIIA-mediated repression of cell cycle genes, including cyclin D1 (CCND1), and the matrix metalloprotease MMP9, respectively (Londhe & Davie, 2011; Nozell et al., 2004). Accordingly, we see both CCND1 and MMP9 are significantly downregulated in NRMT1 KO C2C12 cells (**Figs. 2f, 3f**). As MMP9 is necessary for myoblast migration and engraftment into myofibers (Morgan et al., 2010), these data further suggest that loss of NRMT1 in this model is not preventing cell cycle withdrawal but rather myoblast incorporation into myotubes.

**Fig. 3.**
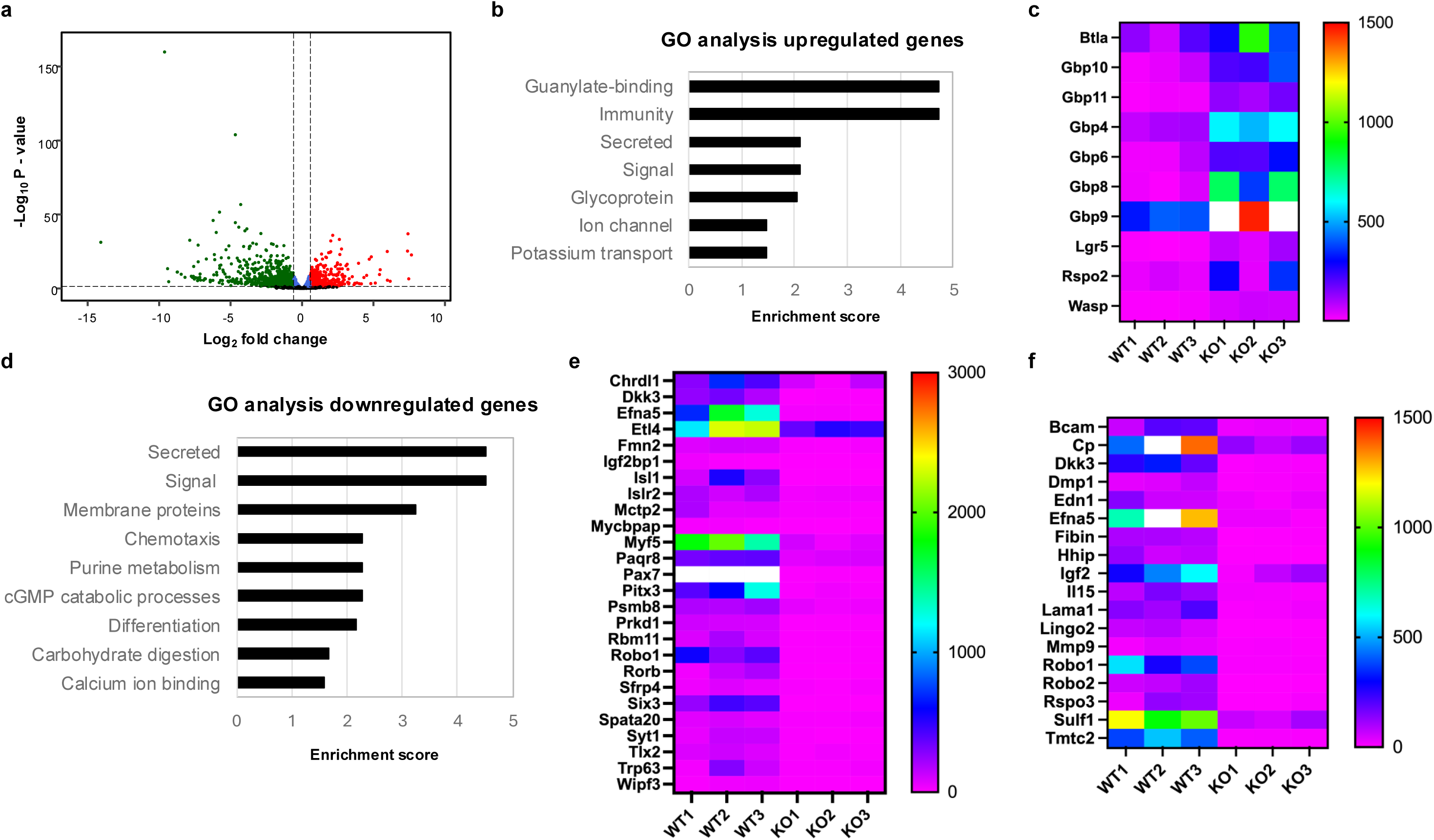
KO day 0 compared to WT day 0. **a** Volcano plot showing genes differentially expressed between NRMT1 knockout (KO) C2C12 cells and wild type (WT) cells at day 0. Blue dots = p-value < 0.05 & Log_2_FC ≤ Log_2_ (1.5); green dots = p-value < 0.05 & Log_2_FC < -Log_2_(1.5), red dots = p-value < 0.05 & Log_2_FC > Log_2_(1.5); black dots = p-value ≥ 0.05. **b** GO analysis of the 101 genes upregulated greater than two-fold in KO cells. **c** Heatmap of upregulated genes with signal peptides or involved in interferon signaling. **d** GO analysis of the 309 genes downregulated greater than two-fold in KO cells. Heatmaps of downregulated genes that play a role in (**e**) differentiation or chemotaxis or (**f**) contain signal peptides. Scale bars on heatmaps denote normalized counts. White bars indicate values are out of range.

We also found 309 genes significantly downregulated greater than twofold in the KO cells (**Fig. 3a**). These included genes encoding for secreted and membrane proteins and genes involved in chemotaxis and differentiation (**Fig. 3d-f**). In addition to *Pax7* and *Myf5,* many of these genes also play roles in myogenic differentiation and migration (**Figs. 3e,f**). R-spondin 3 (RSPO3), is a driver of myogenesis known to increase in C2C12 cells after one day in differentiation media, and its knockdown results in decreased *Myf5* expression, myogenic differentiation, and myotube formation (Han et al., 2011). Insulin-like growth factor 2 (IGF2), and its translational activator insulin-like growth factor 2 binding protein 1 (IGF2BP1), are also critical for muscle growth (Florini et al., 1991; Luo et al., 2024), as IGF2 serves as an autocrine differentiation-promoting factor that is necessary for terminal differentiation of myoblasts (Florini et al., 1991). Mutation of the transcription factor paired-like homeodomain 3 (PITX3) results in withdrawal from the cell cycle and premature differentiation of satellite cells (L’Honoré et al., 2018), and loss of fibin in C2C12 cells inhibits their differentiation into myotubes (Wang et al., 2024).

Ephrin A5 (EFNA5) helps regulate myotube formation. It is expressed on interstitial cells, and through NF-κB-mediated signaling, promotes myoblast migration to the tips of myofibers (Gu et al., 2016). As mentioned above, the metalloprotease MMP9 is also necessary for myoblast engraftment (Morgan et al., 2010), as is its activator DMP1 (Karadag et al., 2005), both of which are found downregulated in the NRMT1 KO cells at day 0 as compared to WT cells (**Fig. 3f**). In addition, there are a number of downregulated genes known to be involved in other types of stem cell chemotaxis (**Fig. 3e,f**). Dickkopf-3 (DKK3) functions in vascular progenitor cell migration (Issa Bhaloo et al., 2018), and the vasoconstrictive peptide endothelin-1 (EDN1/ET-1) serves as a chemoattractant for blood monocytes and neutrophils (Achmad & Rao, 1992; Hofman et al., 1998). Roundabout 1 (ROBO1) functions in migration of neural progenitor cells (Guerrero-Cazares et al., 2017), and roundabout 2 (ROBO2) functions in migration of hepatic stellate cells (Zeng et al., 2018). These data suggest that while WT cells are primed for chemotaxis signaling under basal growth conditions, NRMT1 KO cells may not be able to initiate the chemotaxic signaling necessary for myotube formation.

### Differences in expression after one day of differentiation

After one day of growth in differentiation media, we found 261 genes were significantly upregulated greater than two-fold in the NRMT1 KO line as compared to WT (**Fig. 4a**). In addition to *MyoG* (*MyoD* again did not meet the two-fold cut off), these included genes that encode motor proteins and cytoskeletal proteins, and genes involved in muscle contraction and the troponin complex (**Fig. 4b**). Interestingly, this panel of genes was almost identical to the muscle-specific genes found significantly upregulated in WT cells when comparing between day 0 and day 1 (**Figs. 1d, 4c**). However, the KO cells express them significantly higher than WT cells after one day of differentiation (**Fig. 4c**), indicating they may be trying to compensate for a compromised muscle differentiation program.

**Fig. 4.**
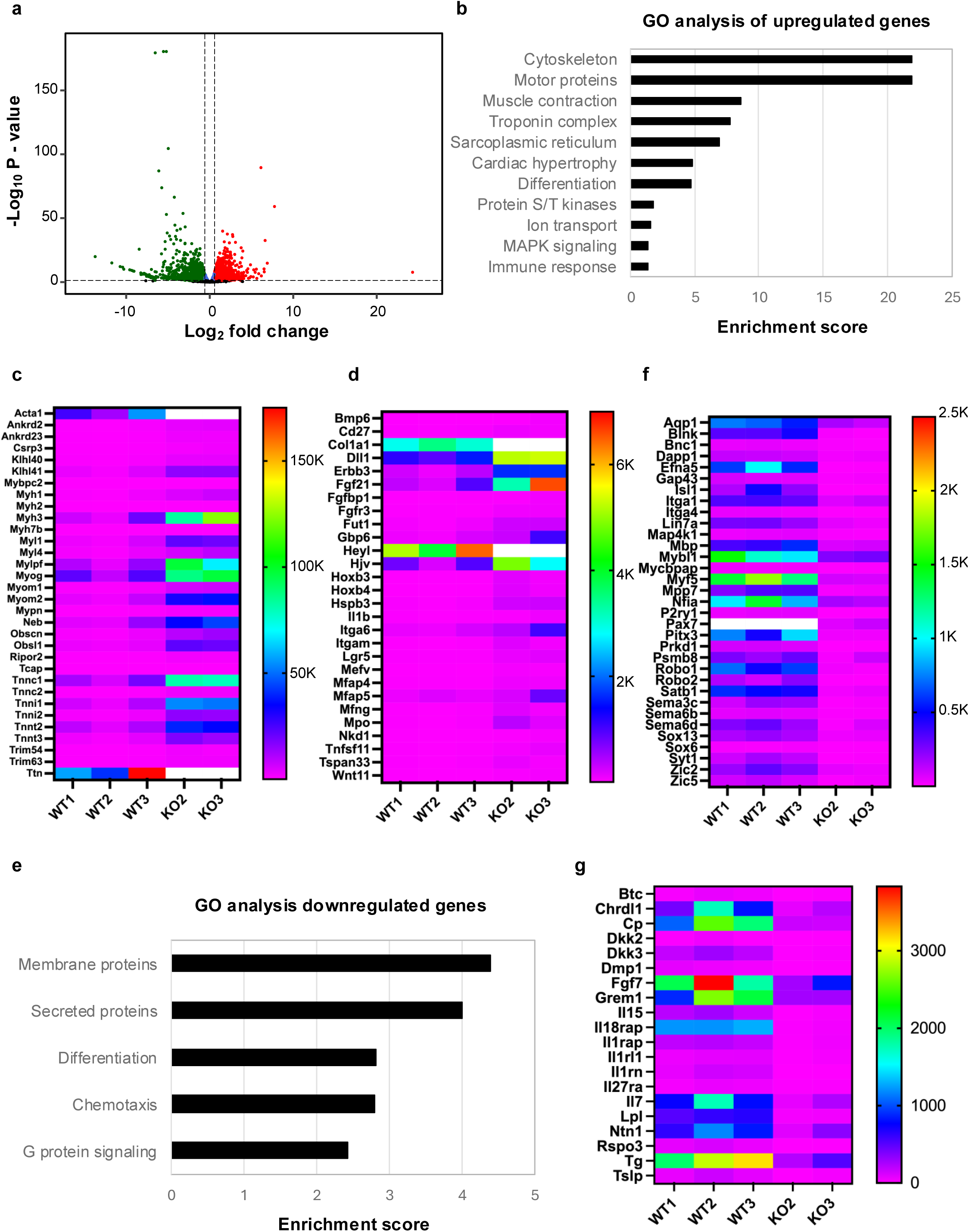
KO day 1 compared to WT day 1. **a** Volcano plot showing genes differentially expressed between NRMT1 knockout (KO) C2C12 cells and wild type (WT) cells at day 1. Blue dots = p-value < 0.05 & Log_2_FC ≤ Log_2_ (1.5); green dots = p-value < 0.05 & Log_2_FC < -Log_2_(1.5), red dots = p-value < 0.05 & Log_2_FC > Log_2_(1.5); black dots = p-value ≥ 0.05. **b** GO analysis of the 261 genes upregulated greater than two-fold in KO cells. Heatmaps of (**c**) muscle-specific transcripts and (**d**) genes involved in differentiation upregulated at day 1. **e** GO analysis of the 486 genes downregulated greater than two-fold in KO cells. Heatmaps of downregulated genes that play a role in (**f**) differentiation or chemotaxis and (**g**) secreted proteins. Scale bars on heatmaps denote normalized counts. White bars indicate values are out of range.

One contributing factor to this hindered differentiation could be that, along with muscle-specific transcripts, we see that KO cells also upregulate genes involved in the differentiation of other lineages (**Fig. 4d**). Specifically, a group of genes that clusters around hemojuvelin (HJV/HFE2) encodes proteins involved in bone differentiation, including BMP6, COL1A1, ERBB3, FGF21, FGFBP1, FGFR3, and TNFSF11/RANKL (Dacic et al., 2001; Dambroise et al., 2020; Hoppman et al., 2010; Lin et al., 2008; Wei et al., 2012; Zhang et al., 2010; Zhao, 2018) (**Fig. 5b**). Another group of genes that clusters around myeloperoxidase (MPO) encodes proteins involved in myeloid differentiation, including CD27, IL-1β, ITGA6, ITGAM, and MEFV (Centola et al., 2000; Nauseef et al., 1988; Nolte et al., 2005; Phillips et al., 2018; Pietras et al., 2016; Ripperger et al., 2015) (**Fig. 5b**). Neither of these clusters are upregulated in WT cells at day 1, which show a striking absence of other lineages branching from the main muscle cluster as compared to the KO cells (**Fig. 5a,b**). These data indicate that while NRMT1 KO cells can initiate the muscle differentiation program, unlike WT cells, they cannot do this exclusively.

**Fig. 5.**
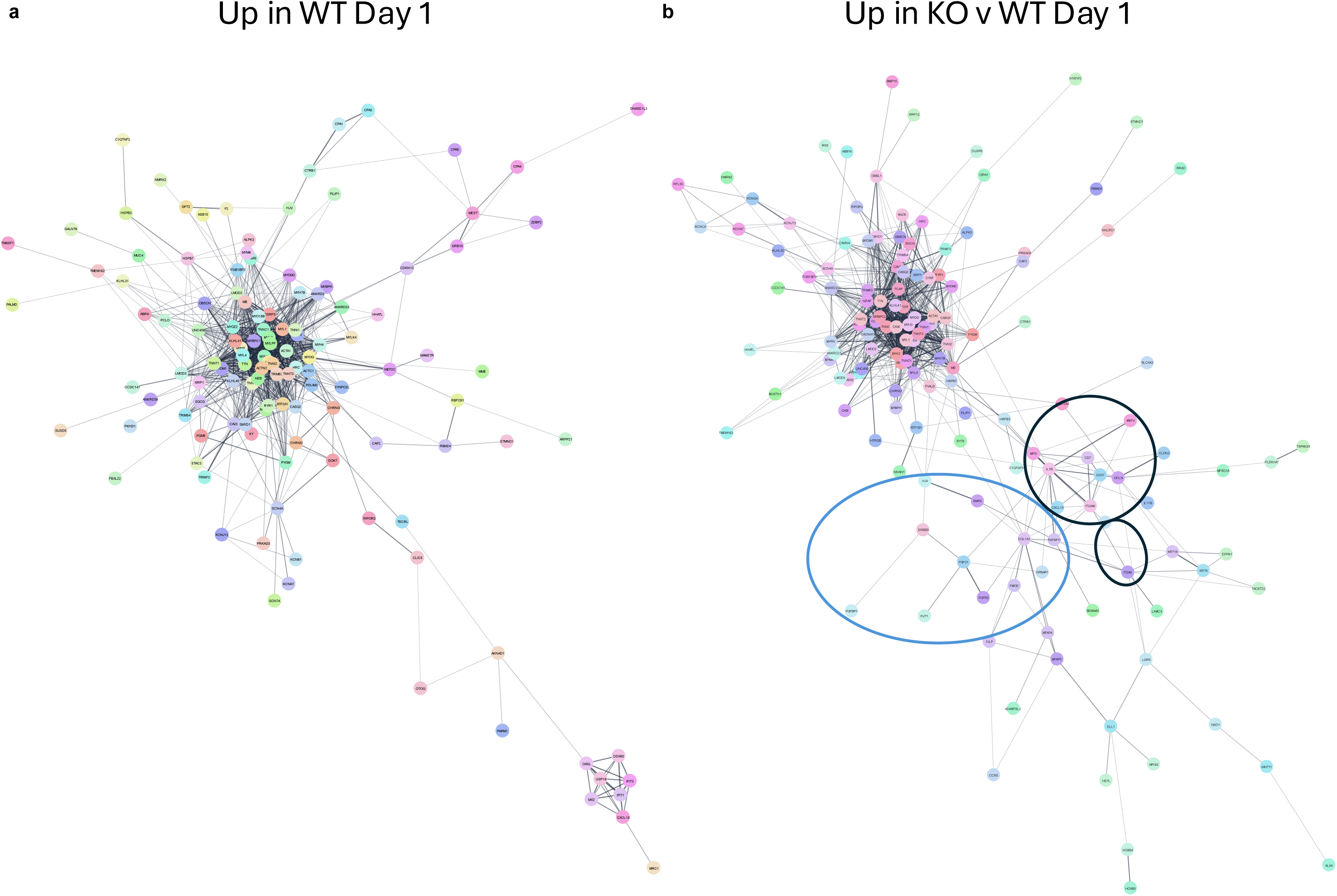
Interaction networks showing major clusters of genes with differential expression at day 1. **a** Genes upregulated in WT cells at day1 as compared to WT cells at day 0. **b** Genes upregulated in KO cells at day 1 as compared to WT cells at day 1. The hemojuvelin (HJV) cluster is circled in blue, and the myeloperoxidase (MPO) clusters are circled in black.

We also found 486 genes significantly downregulated more than two-fold in the KO cells after one day of differentiation (**Fig. 4a**). These included genes that encode for plasma membrane and secreted proteins and genes that encode for proteins involved in differentiation and chemotaxis (**Fig. 4e**). Many of the downregulated membrane and secreted proteins have known roles in muscle differentiation and cluster together in interaction networks (**Fig. 4f,g; Sup. Fig. 2**). Fibroblast growth factor 7 (FGF7) promotes satellite cell proliferation (Ma et al., 2024), and a variety of genes that cluster with it, including *Il15*, *Il18rap*, *Il1rl1*, *Il7, Itga1*, and *Itga4* (**Sup. Fig. 2**), are involved in satellite cell differentiation or migration (Ashraf et al., 2023; Haugen et al., 2010; Ho et al., 2022; Quinn et al., 1997; Wang et al., 2020; Zhou et al., 2020). In addition to *Pax7* and *Myf5*, many of the downregulated differentiation genes also have roles in muscle differentiation (**Fig. 4g**). Lin-7 homolog A (LIN7A) is a major regulator of cell polarity (Gruel et al., 2016) and is known to form a complex with MPP7 (Bohl et al., 2007), which plays a role in satellite cell expansion and self-renewal (Li & Fan, 2017). SRY-box transcription factor 6 (SOX6) facilitates skeletal muscle differentiation by regulating the switch between slow and fast skeletal muscle fiber development (Zhang et al., 2022), and zinc finger protein of the cerebellum 2 (ZIC2) is a transcription factor that promotes *Myf5* expression (Pan et al., 2011).

The downregulated genes involved in chemotaxis at day 0 that remain significantly downregulated greater than two-fold at day 1 of differentiation in the KO cells include *Efna5, Dmp1, Dkk3, Robo1,* and *Robo2* (**Figs. 4f, g**). *Mmp9 and Edn1* remain downregulated but less than two-fold (data not shown). In addition, there are a number of newly downregulated chemotaxic genes at day 1 (**Fig. 4f**), most of which currently have few documented roles in skeletal muscle migration. Dual adaptor for phosphotyrosine and 3-phosphoinostitides 1 (DAPP1/BAM32) is an adaptor protein downstream of phosphoinositide 3-kinase (PI3K) that regulates Rac-1-mediated cytoskeletal rearrangement, actin remodeling, and lamellipodia formation (Hao et al., 2020). Along with MMP9, PI3K signaling is necessary for muscle precursor cell migration in the mouse tongue (Choi et al., 2020), so it is possible DAPP1 plays a role in myoblast migration through its role in PI3K signaling. The water channel aquaporin 1 (AQP1) is necessary for endothelial cell migration, as it is hypothesized that water and ion uptake at the tips of lamellipodium could alter local osmotic gradients and the hydrostatic pressure necessary for cell protrusions (Verkman, 2005). Here, it is possible AQP1 could normally function to regulate myoblast protrusions during directed migration to myotubes/myofibers. Semaphorins are proteins known to direct both axon guidance and cancer metastasis. While semaphorins 3C and 6B (SEMA3C and 6B) seem to follow these traditional roles (Andermatt et al., 2014; Ferrario et al., 2012; Li et al., 2021; Tam et al., 2017), semaphorin 6D (SEMA6D) has an additional role in cardiac morphogenesis (Toyofuku et al., 2004). SEMA6D can exert migration-promoting activity or migration-inhibitory activity on cardiac explants, depending on their source (Toyofuku et al., 2004). Its unique role in cardiac morphogenesis suggests it could also be working here as a chemotactic signal in skeletal muscle. These data indicate NRMT1 loss inhibits complete differentiation into muscle fibers by prohibiting activation of a subset of muscle-specific differentiation pathways (though not all) and blocking myoblast migration into myotubes/myofibers.

## Discussion

Taken together, these RNA-sequencing results reveal several interesting consequences of NRMT1 loss during myoblast growth and differentiation. First, though NRMT KO C2C12 cells exhibit significantly decreased *Pax7* and *Myf5* expression, they are able to still express *MyoD* and *MyoG* and initiate a muscle-specific differentiation program very similar to WT cells (**Figs. 1d, 4c**). Second, though loss of NRMT1 has previously been shown to result in decreased levels and function of RB in NSCs (Catlin et al., 2021), NRMT1 KO myoblasts are able to significantly downregulate expression of genes involved in the cell cycle and DNA replication (**Figs. 2f,g**), which corresponds with our previous data showing NRMT1 loss decreases the growth rate of C2C12 cells (Tooley et al., 2021) and suggests an inability to exit the cell cycle is not blocking their terminal differentiation into myotubes. Finally, these results confirm that NRMT1 KO C2C12 cells fail to completely repress differentiation programs from different lineages (**Fig. 4d**), and this, combined with altered transcription of the signaling molecules necessary to promote myoblast migration (**Fig. 4g,f**), could be driving the observed terminal differentiation defects.

Though *MyoD* transcription is thought to be mainly activated through the action of PAX7 and MYF5, it can alternatively occur through the action of other transcription factors such as SIX1/4 and FOXO3 (Hernández-Hernández et al., 2017; Hu et al., 2008). FOXO3 seems to work cooperatively with PAX7, as neither can form a stable DNA-activating complex alone (Hu et al., 2008). However, SIX1 and SIX4 seem to work together to promote *MyoD* expression independently of PAX7 (Liu et al., 2013; Santolini et al., 2016). MYOD can act as its own transcriptional activator (Thayer et al., 1989), and SIX1 is necessary for MYOD to access its own enhancer (Liu et al., 2013). It has been proposed that SIX1 works to open the chromatin at the core enhancer region upstream of the *MyoD* transcription start site and facilitate MYOD recruitment (Liu et al., 2013). As both SIX1 and SIX4 maintain comparable expression between KO and WT cells at both time points (data not shown), it is possible that they are responsible for maintaining *MyoD* expression in the absence of PAX7 and MYF5. MYOD is a transcriptional activator of *MyoG* upon stimulation of differentiation (Deato et al., 2008), and its presence is likely responsible for the observed increase of *MyoG* expression and initiation of the muscle-specific transcriptional program in the KO cells. The expression patterns seen here for *MyoD* and *MyoG* differ from our previous qRT-PCR data, where neither were detectable in the KO cells at day 0 and no increase was seen in *MyoG* at day 1 (Tooley et al., 2021). We attribute this to the increased sensitivity of RNA-sequencing and the optimization of cell confluency conditions during differentiation.

Besides an inability to initiate the muscle-specific transcriptional program, another potential contributor to the inability of NRMT1 KO cells to form myotubes (Tooley et al., 2021) was failure to exit the cell cycle. In NSCs, we have previously shown that levels of phosphorylated RB, a NRMT1 substrate, are increased upon NRMT1 KO, which results in decreased overall levels of RB and an increase in RB-regulated cell cycle transcripts, including cyclins A2 (CCNA2) and E2 (CCNE2) (Catlin et al., 2021). This corresponds with an inability of neurons to permanently exit the cell cycle and undergo terminal differentiation (Catlin et al., 2021). The terminal differentiation phenotypes occurring in myoblasts appear to be mechanistically different than those in NSCs, as NRMT1 KO C2C12 downregulate a number of cell cycle and DNA replication genes, including CCNA2 and CCNE2 (**Fig. 2f,g**). Another striking observation in the KO cells is their massive downregulation of histone gene expression (**Fig. 2h**). This is traditionally seen at the end of S phase, after DNA replication has been completed, and at the initiation of differentiation (Mei et al., 2017; Stein et al., 1996). These data suggest that the KO cells are withdrawing from the cell cycle and support our previous data showing the KO cells exhibit slowed proliferation (Tooley et al., 2021).

Our previous data also showed that basally growing NRMT1 C2C12 KO cells expressed markers associated with osteoblastic differentiation (Tooley et al., 2021). Results from these RNA-sequencing experiments confirm this and suggest it may be due to altered Wnt/β-catenin signaling pathways. Though expression of most *Wnt* genes is not statistically different between the two cell lines at either time point, *Wnt11* expression significantly increases after day 1 of differentiation in the KO cells (**Fig. 4d**). WNT11 is an important signaling molecule for osteoblast maturation (Friedman et al., 2009). The Wnt signaling agonist RSPO2 and its receptor leucine-rich repeat-containing G-protein coupled receptor 5 (LGR5) are also upregulated in the KO cells (**Fig. 3c**), and both have additionally been shown to promote osteoblastic differentiation (Friedman et al., 2009; Yu et al., 2021). In contrast, a number of Wnt signaling antagonists were shown to be downregulated in the KO cells (**Figs. 3e, 4g**), including DKK2, DKK3, SIX3, and SFRP4 (Ford et al., 2013; Hirata et al., 2009; Lagutin et al., 2003; Martin Flores et al., 2024). This altered Wnt signaling could also lead to the observed upregulation of genes involved in myeloid differentiation, as constitutively active β-catenin can cause activation of myeloid markers in committed lymphoid progenitors (Baba et al., 2006). It has been proposed that the diminished lineage restriction seen with β-catenin activation could be due to significant epigenetic changes (Malhotra & Kincade, 2009), which given the observed, massive downregulation of histone genes in KO cells (**Fig. 2h**), could also be occurring with NRMT1 loss.

Finally, we had previously seen that NRMT1 KO C2C12 cells were unable to form myotubes in culture (Tooley et al., 2021). This could be a direct effect, mediated through a physical inability of the cells to migrate. Many myosin light chains are direct targets or NRMT1 (Tooley et al., 2010), and transfection of NIH 3T3 cells with a MYL9 construct that cannot be N-terminally methylated promotes significant cell spreading/adhesion (Nevitt et al., 2018), which could inhibit migration. The inability to form myotubes could also be an indirect result of disrupted chemotaxis, mediated through alteration of secreted protein levels. The trafficking of secreted proteins is primarily regulated through N-terminal signal sequences (Martoglio, 2003). Processing of these signal sequences, including co- or post-translational modification and cleavage, helps direct a protein through the secretory process (Martoglio, 2003; Rutkowski et al., 2003). One known peptidase of secreted proteins, dipeptidyl peptidase 9 (DPP9), shares consensus sequence requirements with NRMT1 and has a number of overlapping targets with signal sequences (Petkowski et al., 2012; Zhang et al., 2015). It is possible that N-terminal methylation by NRMT1 blocks N-terminal cleavage by DPP9. Loss of NRMT1 would then significantly alter composition of the secretome, which could result in compensatory alteration of transcript levels and the misregulation of chemotaxic gene expression seen in the KO cells.

These RNA-sequencing experiments have shed some light on how NRMT1 loss prevents myotube formation of C2C12 myoblasts. They indicate that the KO cells can differentiate down muscle-specific lineages and withdraw from the cell cycle. However, the KO cells are not able to successfully repress the differentiation programs of other lineages and have significant misregulation of secreted protein transcription, including those involved in chemotaxis. Future work will look to determine how NRMT1 loss leads to decreased PAX7 and MYF5 expression, as there are a number of transcription factors and modulators of chromatin structure that are direct targets of NRMT1(Conner et al., 2022; Dai et al., 2013; Sathyan et al., 2017; Tooley et al., 2010). We will also try to discern whether loss of NRMT1 directly impacts cell migration through impaired cytoskeletal rearrangement or indirectly impacts cell migration through alteration of the secretome. Comparing how NRMT1 regulates myoblast differentiation with how it affects NSC differentiation will help determine if the premature aging phenotypes seen in *Nrmt1^-/-^*mice are due to general defects in stem cell maintenance or if there are tissue-specific consequences.

## Acknowledgments and Funding Information

This work was supported by a research grant from the National Institutes of Health to CST [GM144111].

---

**Sup. Fig. 1.**
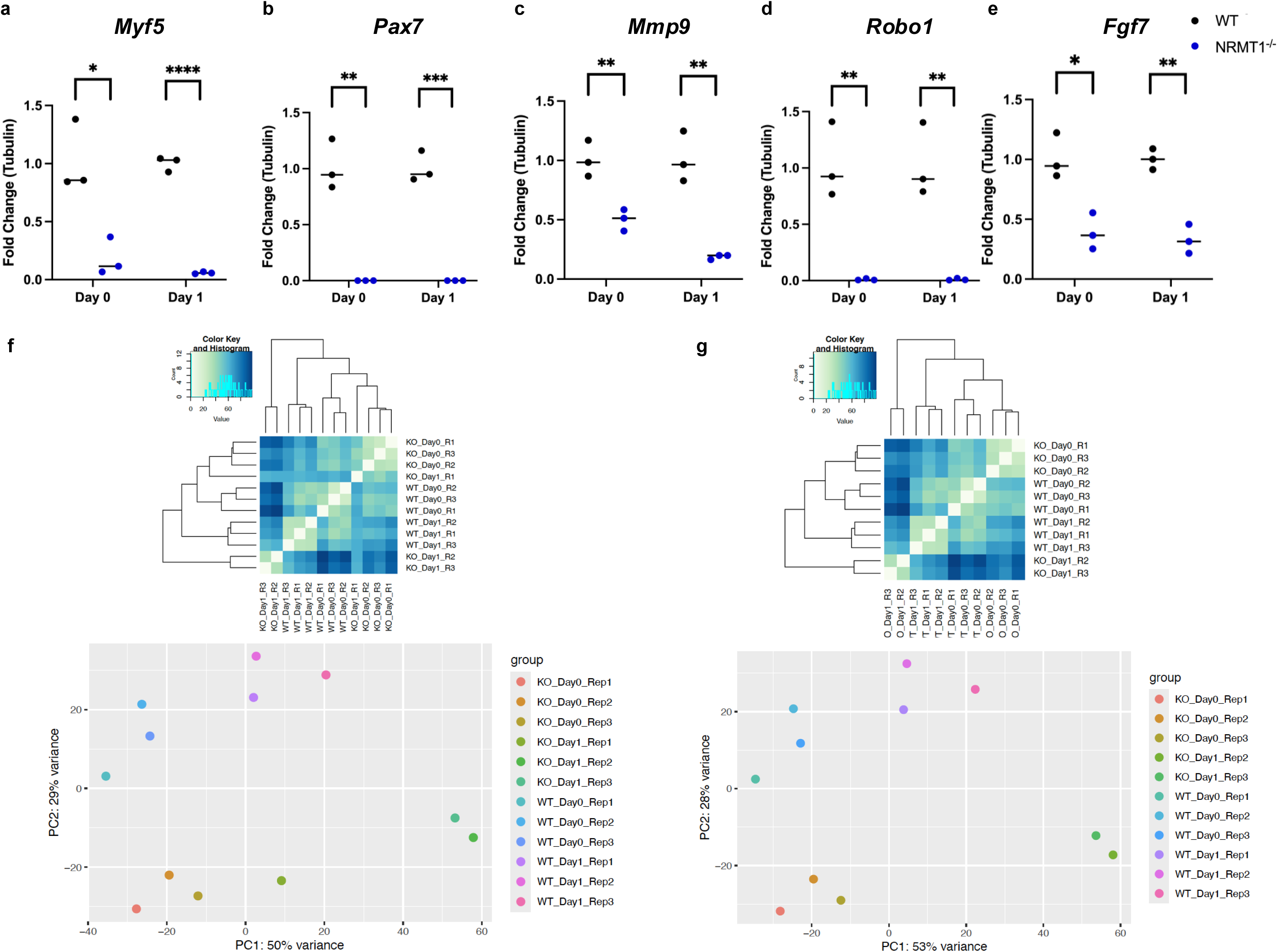
Quality control analysis. **a-e** Quantitative PCR verification of RNA-sequencing results. **f** Hierarchical clustering was performed on all twelve gene samples and the results showed that the KO Day1 Rep1 sample (olive green) was a significant outlier and needed to be removed. **g** Hierarchical clustering performed with KO Day1 Rep1 removed. These samples were used for all further analysis.

**Sup. Fig. 2.**
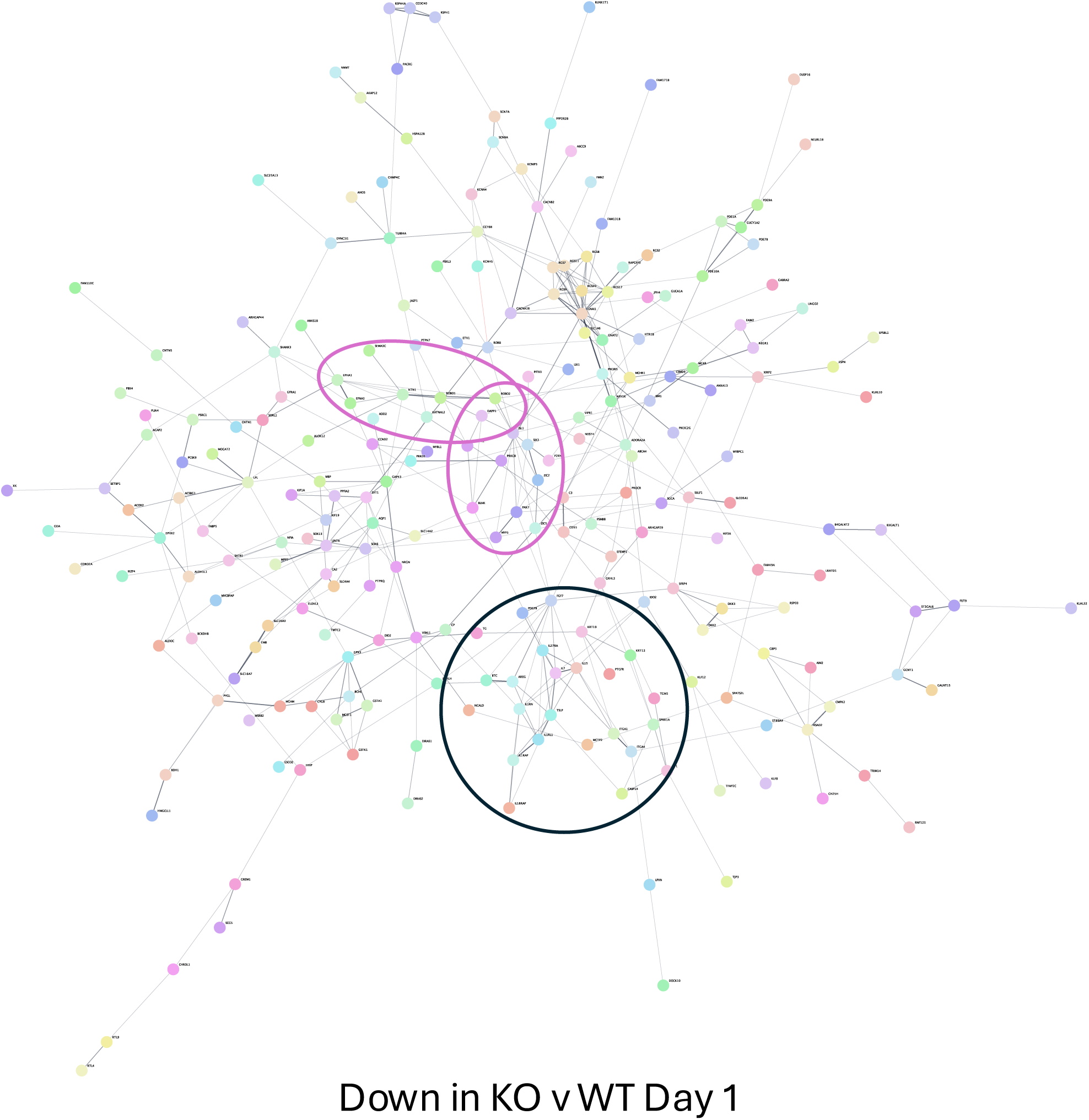
Interaction network of genes downregulated in KO cells at day 1 as compared to WT cells at day 1. The ROBO2 clusters, which include ROBO1, EFNA5, SEMA3C, DAPP1, PAX7, MYF5, and ZIC2, are circled in purple. The FGF7 cluster is circled in black.

